# Subtype-Specific SASP Dynamics Predict Mesenchymal Transition in Recurrent Glioblastoma

**DOI:** 10.64898/2026.01.15.699753

**Authors:** Varshith Kotagiri, Ava Teng

## Abstract

**Background:** Therapy-induced senescence is hypothesized to promote glioblastoma recurrence through senescence-associated secretory phenotype (SASP) factors, yet no longitudinal study has tracked SASP dynamics across matched primary-recurrent pairs. We used the GLASS consortium to characterize ΔSASP trajectories and their clinical implications.

**Methods:** We analyzed 167 matched primary-recurrent GBM pairs from the GLASS consortium. SASP scores were calculated from a 10-gene signature. Patients were classified by ΔSASP trajectory (Accumulators vs Clearers). We tested associations with proneural-to-mesenchymal transition (PMT) and validated findings in TCGA (n=52).

**Results:** SASP dynamics were subtype-specific: Mesenchymal tumors showed significant SASP clearance (ΔSASP = −0.83, p = 0.0001), while Classical tumors remained stable (p = 0.86). Among individual genes, HGF was the only factor that increased at recurrence (Δ = +0.23, p = 0.016), while IL1A, IL1B, and MMP9 decreased significantly. ΔSASP strongly predicted mesenchymal transition (OR = 5.06, 95% CI: 2.30-11.18, p < 0.0001), which itself conferred worse survival (HR = 2.48, p = 0.0001). Strikingly, Clearers were nearly completely protected from PMT (2.4% vs 27.3% in Accumulators, p = 0.0008). ΔSASP correlated with immunosuppressive infiltration (ρ = 0.65 with M2 macrophages, p < 0.0001). Baseline SASP predicted worse survival (HR = 1.65, p < 0.0001), an effect that persisted in IDH-wildtype tumors (HR = 1.32, p = 0.015) and was validated in the CGGA cohort (HR = 1.35, p = 0.003). External validation confirmed the SASP-Mesenchymal association in both TCGA (d = 1.75) and CGGA (d = 1.14).

**Conclusions:** SASP trajectories are heterogeneous and subtype-specific. Patients who fail to clear SASP represent a high-risk subgroup undergoing mesenchymal transition and immunosuppression. These Accumulators may be optimal candidates for senolytic therapy, while Clearers demonstrate effective senescence surveillance that could be therapeutically reinforced.

## Introduction

Glioblastoma (GBM) is the most common and lethal primary brain tumor in adults, with an annual incidence of approximately 3.2 per 100,000 and a median overall survival of only 14–16 months despite aggressive multimodal therapy. The current standard of care, maximal safe surgical resection followed by concurrent temozolomide and radiotherapy (the Stupp protocol), has remained largely unchanged for nearly two decades^[1,2]^. Despite initial treatment response, recurrence is virtually universal. The median time from diagnosis to first recurrence is 6–9 months, and salvage therapies offer limited benefit. Understanding the molecular mechanisms that drive recurrence is therefore critical for developing more effective therapeutic strategies.

Cellular senescence is a state of permanent cell cycle arrest coupled with profound secretory reprogramming. Senescent cells remain metabolically active and secrete a complex mixture of cytokines (IL-6, IL-1β), chemokines (CCL2, CXCL8), growth factors (HGF, VEGF), and matrix metalloproteinases collectively termed the senescence-associated secretory phenotype (SASP)^[3,4]^. Both temozolomide and ionizing radiation are potent inducers of senescence. Studies have demonstrated that temozolomide-induced senescence occurs at approximately four-fold higher levels than apoptosis in GBM cells, with greater than 80% of tumor cells surviving initial chemotherapy entering a senescent state^[5,6]^. This creates a therapeutic paradox: while senescence acutely suppresses tumor proliferation, the chronic accumulation of SASP factors may promote survival and proliferation of non-senescent tumor cells, ultimately driving recurrence^[7]^. Recent work by Salam et al. demonstrated that senescent cells, comprising less than 7% of tumor mass, are sufficient to promote GBM progression, and that higher senescence scores correlate with shorter patient survival^[8]^.

The prevailing model predicts that senescence burden should increase from primary to recurrent tumors as therapy-induced senescent cells accumulate. Higher ΔSASP should therefore correlate with faster and more aggressive recurrence. However, this hypothesis has never been tested with longitudinal matched-pair data. A recent comprehensive review explicitly identified this knowledge gap, noting that “few studies to date have focused on glioma, likely due to the absence of available longitudinal tissue sets” and that “it remains unknown how this relative senescent burden at baseline is impacted after standard glioma therapy based on longitudinal samples”^[9]^.

Transcriptional subtyping has identified distinct GBM molecular subtypes—Proneural, Classical, Mesenchymal, and Neural—with the Mesenchymal subtype associated with worst prognosis and highest inflammatory signature^[10]^. Single-cell RNA sequencing has further refined this classification, revealing four cellular states (NPC-like, OPC-like, AC-like, and MES-like) that coexist within individual tumors, with NF1 mutations favoring the mesenchymal-like state^[11]^. Importantly, subtype is not fixed: approximately 50% of recurrent GBMs acquire a Mesenchymal phenotype regardless of initial classification, a phenomenon termed proneural-to-mesenchymal transition (PMT)^[12]^. PMT is driven by NF-κB activation, which notably also serves as the master transcriptional regulator of SASP^[13,14]^. This mechanistic overlap suggests a potential link between SASP accumulation and mesenchymal transition that has not been quantitatively investigated.

Prior studies of senescence in GBM have relied on cross-sectional data, analyzing senescence signatures at a single timepoint^[8]^. While these studies have established that higher senescence burden correlates with worse prognosis, they cannot assess the dynamic change in SASP from primary to recurrent tumors within individual patients. The GLASS (Glioma Longitudinal Analysis) Consortium provides a unique resource: matched RNA-sequencing data from primary and recurrent tumors in the same patients^[15]^. This enables, for the first time, calculation of ΔSASP—the change in senescence burden across the treatment interval—and classification of patients by their senescence trajectory.

### Novel contributions of this study

1. First longitudinal ΔSASP analysis in matched GBM pairs. 2. Discovery of subtype-specific SASP dynamics. 3. Quantitative link between ΔSASP and PMT. 4. Trajectory-based patient stratification with therapeutic implications.

## 2. Methods

### 2.1 Study Design and Cohorts

#### Discovery Cohort: GLASS Consortium

We analyzed the GLASS consortium 2022 data release, which provides matched primary-recurrent RNA-sequencing for glioma patients^[15,16]^. Inclusion criteria were: (1) histologically confirmed glioblastoma, (2) RNA-seq data available for both primary and first recurrent tumor, and (3) transcriptional subtype annotation for both timepoints. The final analysis cohort comprised 167 patients with complete matched pairs. IDH status was available for 155 patients (92.8%): 118 (70.7%) IDH-wildtype, 37 (22.2%) IDH-mutant, and 12 (7.2%) unknown. Primary tumor subtypes were Classical (n=145, 86.8%) and Mesenchymal (n=22, 13.2%).

#### Validation Cohort

**TCGA-GBM**. For external validation, we analyzed The Cancer Genome Atlas GBM cohort accessed via cBioPortal^[17]^. This cross-sectional cohort (n=52 with complete data) was used to validate SASP-subtype associations identified in GLASS.

#### External Validation Cohorts

Two independent cohorts were used for external validation. The Cancer Genome Atlas (TCGA) GBM cohort (n=52) provided cross-sectional validation of SASP-subtype associations. The Chinese Glioma Genome Atlas (CGGA) cohort (n=225 GBM samples) was used to validate SASP-survival associations and SASP-M2 correlations. For CGGA, molecular subtypes were predicted using single-sample gene set enrichment analysis with Verhaak 2010 classifier gene sets; 89.8% of samples were classified with high confidence.

### 2.2 SASP Signature and Score Calculation

The SASP signature comprised 10 canonical SASP genes available in both datasets: IL6, IL1A, IL1B, CCL2, CXCL1, HGF, VEGFA, MMP3, MMP9, SERPINE1. CXCL8 (IL-8) was excluded due to unavailability in the expression matrix. Gene selection was based on established SASP literature^[4]^. SASP scores were calculated as follows: 1. Log2-transform TPM values: log2(TPM + 1). 2. Z-score normalizes each gene across all samples in the cohort. 3. SASP score = mean of z-scored SASP genes per sample. 4. ΔSASP = SASP(recurrent) − SASP(primary). Positive ΔSASP indicates SASP accumulation; negative ΔSASP indicates SASP clearance.

### 2.3 Trajectory Classification

Patients were classified into trajectory groups using tertile-based cutoffs:

| Trajectory | Definition | N (%) |
| --- | --- | --- |
| Accumulator | $\Delta\text{SASP} > +0.135$ (67th percentile) | 56 (33.5%) |
| Stable | $-0.333$ to $+0.135$ | 55 (32.9%) |
| Clearer | $\Delta\text{SASP} < -0.333$ (33rd percentile) | 56 (33.5%) |

### 2.4 Subtype and Transition Analysis

Transcriptional subtypes were obtained from GLASS annotations based on the Verhaak classification system^[10]^. Proneural-to-Mesenchymal Transition (PMT) was defined as any non-Mesenchymal primary tumor that became Mesenchymal at recurrence. Association between ΔSASP and PMT was tested using logistic regression, with results reported as odds ratios (OR) with 95% confidence intervals.

### 2.5 Immune Deconvolution

Immune cell fractions were estimated using marker gene signature scoring for M2 macrophages, monocytes, and CD8+ T cells. Change in immune infiltration was calculated as Δimmune = fraction(recurrent) − fraction(primary). Correlations between ΔSASP and Δimmune were assessed using Spearman’s rank correlation.

### 2.6 Survival Analysis

Overall survival was measured from date of initial diagnosis to death or last follow-up. Kaplan-Meier curves were generated for trajectory groups and compared using the log-rank test. Cox proportional hazards regression was used to assess the association between baseline SASP score and survival, with results reported as hazard ratios (HR) with 95% confidence intervals.

### 2.7 Statistical Analysis

Continuous variables were summarized as median (IQR) and compared using Wilcoxon rank-sum or Kruskal-Wallis tests. Categorical variables were compared using Chi-square or Fisher’s exact tests. Effect sizes were reported as Cohen’s d for continuous outcomes and odds ratios for binary outcomes. All analyses were performed in Python using pandas, scipy, lifelines, and statsmodels. Statistical significance was defined as two-sided p < 0.05.

### 2.8 Sensitivity Analyses

To assess robustness, SASP-survival associations were tested using three signature definitions: the original 10-gene signature, the Coppé core 6-gene signature (IL6, IL1B, CCL2, MMP3, MMP9, SERPINE1), and an expanded 13-gene signature. To confirm SASP was distinct from generic inflammation, we calculated an inflammation score using non-SASP genes (CRP, TNF, IFNG, CD3E, CD8A) and tested whether SASP retained prognostic significance after adjustment in multivariate Cox regression

## 3. Results

### Cohort Characteristics

The final analysis cohort comprised 167 patients with matched primary-recurrent tumor pairs (Table 1). The cohort was predominantly IDH-wildtype (70.7%) and enriched for Classical subtype at primary diagnosis (86.8%). Median overall survival was 24.0 months, with 146 deaths (86.4%) observed during follow-up.

**Table 1:** Patient Characteristics.

| Characteristic | GLASS (n=167) |
| --- | --- |
| IDH-wildtype | 118 (70.7%) |
| IDH-mutant | 37 (22.2%) |
| IDH-unknown | 12 (7.2%) |
| Primary Subtype - Classical | 145 (86.8%) |
| Primary Subtype - Mesenchymal | 22 (13.2%) |
| Recurrent Subtype - Classical | 137 (82.0%) |
| Recurrent Subtype - Mesenchymal | 27 (16.2%) |
| Recurrent Subtype - Proneural | 3 (1.8%) |
| Median OS (months) | 24.0 |
| Deaths | 146 (86.4%) |

**Table 2:** Gene-Specific ΔSASP Analysis.

| Gene | Mean $\Delta$ | 95% CI | p-value | Direction |
| --- | --- | --- | --- | --- |
| HGF | +0.23 | [0.01, 0.44] | 0.016 | ↑ Increased |
| SERPINE1 | +0.15 | [-0.19, 0.50] | 0.43 | — |
| VEGFA | -0.05 | [-0.47, 0.37] | 0.73 | — |
| MMP3 | -0.07 | [-0.21, 0.07] | 0.90 | — |
| CXCL1 | -0.10 | [-0.31, 0.12] | 0.32 | — |
| IL1A | -0.19 | [-0.33, -0.04] | 0.007 | ↓ Decreased |
| IL6 | -0.25 | [-0.52, 0.01] | 0.10 | — |
| IL1B | -0.28 | [-0.55, 0.00] | 0.016 | ↓ Decreased |
| CCL2 | -0.30 | [-0.62, 0.02] | 0.12 | — |
| MMP9 | -0.41 | [-0.81, -0.01] | 0.041 | ↓ Decreased |

**Table 3:** PMT Rate by Trajectory Group.

| Trajectory | N eligible | N PMT | PMT Rate (95% CI) |
| --- | --- | --- | --- |
| Accumulator | 55 | 15 | 27.3% (17.3–40.2%) |
| Stable | 49 | 4 | 8.2% (3.2–19.2%) |
| Clearer | 41 | 1 | 2.4% (0.4–12.6%) |

### SASP Dynamics Are Subtype-Specific

Overall, ΔSASP showed a trend toward decrease at recurrence (mean = −0.085, median = −0.050), but this did not reach statistical significance (Wilcoxon p = 0.15; Figure 1A). Fifty-five percent of patients showed ΔSASP < 0 (Clearers), while 45% showed ΔSASP > 0 (Accumulators). The paired trajectories for individual patients are shown in Figure 1B. However, stratification by primary subtype revealed strikingly divergent patterns (Figure 1C). Mesenchymal tumors showed highly significant SASP clearance (mean ΔSASP = −0.83, p = 0.0001), while Classical tumors showed no significant change (mean ΔSASP = +0.03, p = 0.86). This subtype-specific effect explains the apparent heterogeneity in the pooled analysis.The magnitude of SASP clearance in Mesenchymal tumors was substantial, representing nearly a full standard deviation decrease in SASP score. This finding suggests that Mesenchymal tumors, which have the highest baseline SASP due to constitutive NF-κB activation^[13]^, may engage senescence clearance mechanisms more effectively than other subtypes. Patients were classified into three trajectory groups based on tertile cutoffs (Figure 1D): Accumulators (n=56, 34%), Stable (n=55, 33%), and Clearers (n=56, 34%).

**Figure 1:**
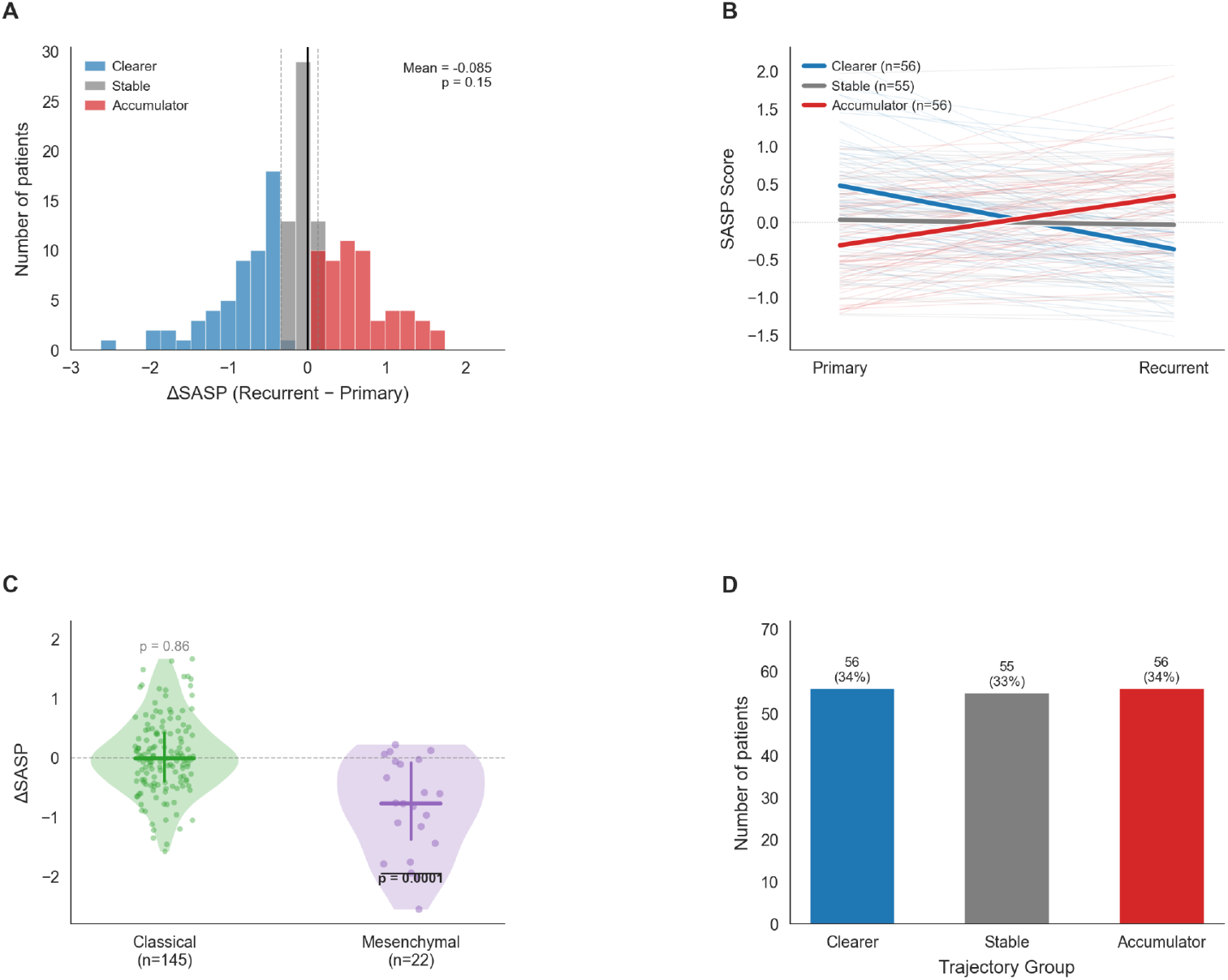
SASP Dynamics at Glioblastoma Recurrence. (A) Distribution of ΔSASP (recurrent minus primary) across 167 matched tumor pairs. Patients are colored by trajectory group:Clearers (blue, ΔSASP < −0.33), Stable (gray), and Accumulators (red, ΔSASP > +0.13). Mean ΔSASP = −0.085, p = 0.15 by one-sample Wilcoxon test. (B) Paired SASP scores showing individual patient trajectories from primary to recurrent tumor. Thick lines represent group means; thin lines represent individual patients. (C) ΔSASP stratified by primary tumor subtype. Mesenchymal tumors (n=22) show significant SASP clearance (mean ΔSASP = −0.83, p = 0.0001), while Classical tumors (n=145) show no significant change (p = 0.86). Violin plots show distribution; horizontal lines indicate median and quartiles. (D) Distribution of patients across trajectory groups based on tertile cutoffs.

### Gene-Specific Analysis: HGF Uniquely Increases

To identify which SASP components drive the observed changes, we analyzed each gene individually (Figure 4). HGF was the only SASP gene that significantly increased at recurrence (Δ = +0.23, 95% CI: 0.01–0.44, p = 0.016). In contrast, several canonical SASP factors decreased significantly: IL1A (Δ = −0.19, p = 0.007), IL1B (Δ = −0.28, p = 0.016), and MMP9 (Δ = −0.41, p = 0.041).

The unique increase in HGF is consistent with the stromal senescence model proposed by Fletcher-Sananikone et al.^[18]^, in which senescent astrocytes in the tumor microenvironment secrete HGF that activates MET signaling in tumor cells. Notably, MET amplification is known to increase at GBM recurrence^[19]^, suggesting a potential HGF/MET paracrine axis that persists despite clearance of tumor-intrinsic SASP. Recent work has demonstrated that SASP-derived HGF directly protects tumor cells from elimination through cell competition mechanisms^[20]^.

### ΔSASP Strongly Predicts Mesenchymal Transition

Among 145 patients with non-Mesenchymal primary tumors, 20 (13.8%) underwent PMT (Figure 2C). Patients who transitioned to Mesenchymal had markedly higher ΔSASP compared to those who maintained their subtype (mean +0.61 vs −0.18, difference = 0.79, Cohen’s d = 1.10; Figure 2A). In logistic regression, ΔSASP was a strong predictor of PMT (OR = 5.06, 95% CI: 2.30–11.18, p < 0.0001). This indicates that each 1-unit increase in ΔSASP was associated with a 5-fold higher odds of mesenchymal transition. When stratified by trajectory group, the protective effect of SASP clearance became strikingly apparent (Figure 2B). Clearers had a PMT rate of only 2.4% (1/41), compared to 8.2% (4/49) in Stable patients and 27.3% (15/55) in Accumulators (Chi-square p = 0.0008). This represents an 11-fold difference in PMT risk between Clearers and Accumulators. Critically, PMT itself was associated with significantly worse survival. Patients who underwent PMT had a 2.5-fold higher hazard of death compared to those who maintained their subtype (HR = 2.48, 95% CI: 1.56–3.94, p = 0.0001), with a median survival of 18.2 months versus 40.1 months (difference = 21.9 months). This establishes the clinical significance of the ΔSASP → PMT axis: Accumulators are not only at higher risk of phenotypic transition, but this transition itself confers substantially worse prognosis.

**Figure 2:**
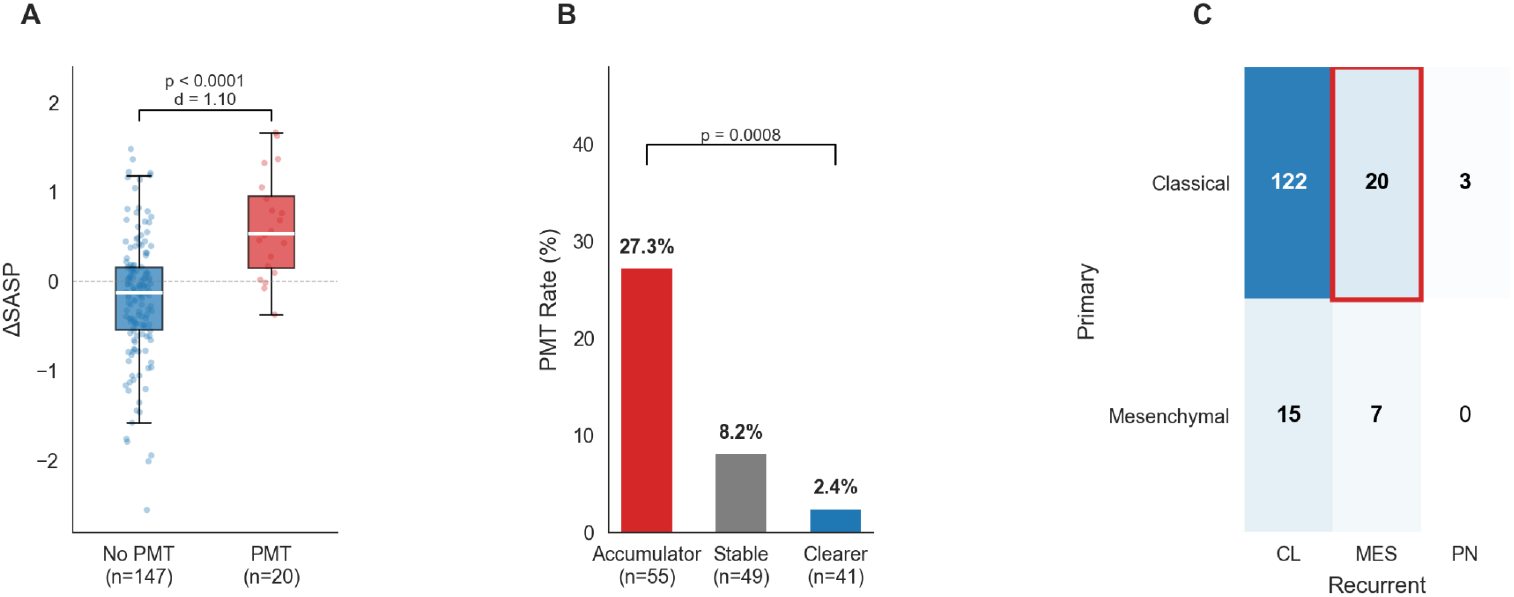
ΔSASP Predicts Proneural-to-Mesenchymal Transition. (A) ΔSASP in patients who underwent PMT (n=20) versus those who did not (n=147). Patients with PMT had significantly higher ΔSASP (mean +0.61 vs −0.18, p < 0.0001, Cohen’s d = 1.10). (B) PMT rate by trajectory group. Accumulators had 27.3% PMT rate compared to 2.4% in Clearers (p = 0.0008). Error bars represent 95% confidence intervals. (C) Subtype transition matrix showing the number of patients transitioning between subtypes from primary to recurrent tumor. Red box highlights PMT events (non-Mesenchymal → Mesenchymal).

### ΔSASP Correlates with Immunosuppressive Infiltration

ΔSASP was strongly correlated with changes in immunosuppressive cell populations at recurrence (Figure 3). The strongest correlation was observed with M2 macrophages (ρ = 0.65, p < 0.0001; Figure 3A), followed by monocytes (ρ = 0.52, p < 0.0001) and CD8+ T cells (ρ = 0.29, p = 0.0002). Accumulators showed increased M2 macrophage infiltration at recurrence, while Clearers showed decreased immunosuppressive infiltration (Figure 3B). This suggests that SASP accumulation promotes an immunosuppressive microenvironment that may contribute to both PMT and treatment resistance.

**Figure 3:**
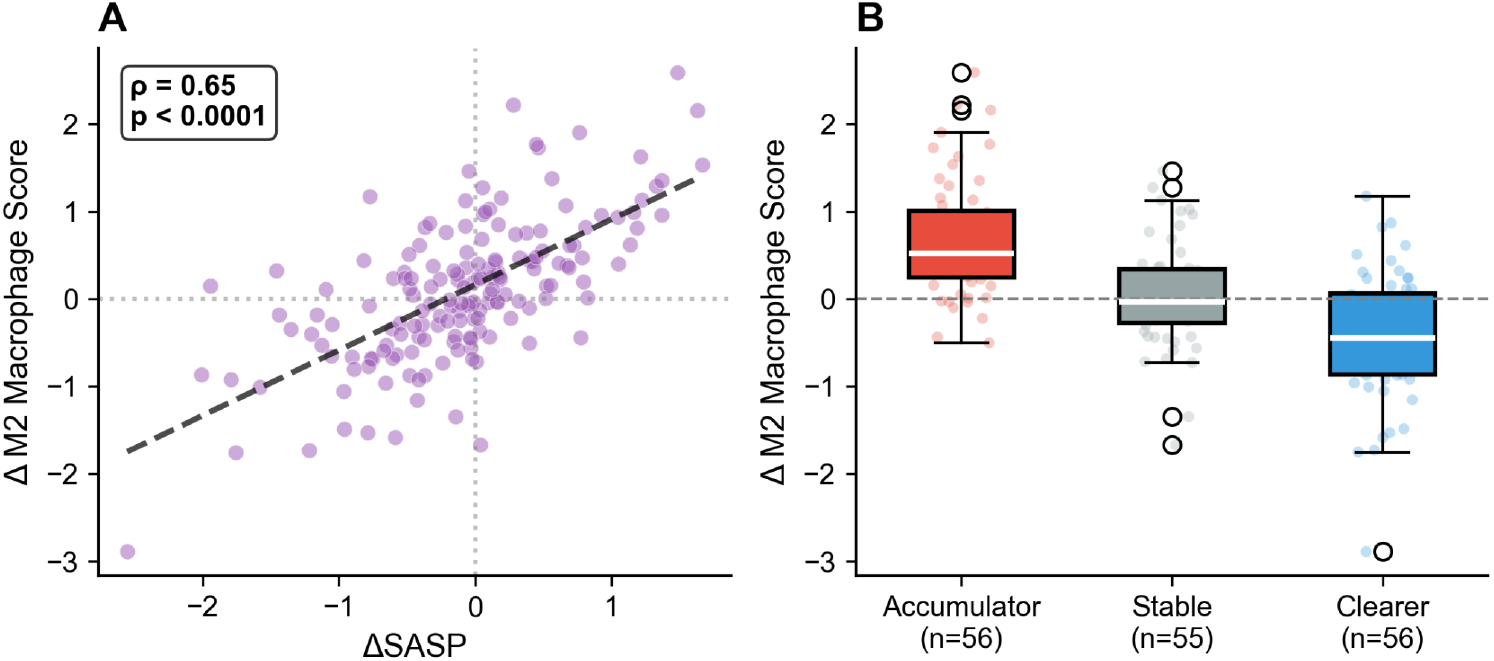
ΔSASP Correlates with Immune Microenvironment Changes. (A)Scatter plot of ΔSASP versus change in M2 macrophage score. Points colored by trajectory group. Spearman ρ = 0.65, p < 0.0001. (B) Change in M2 macrophage score by trajectory group. Accumulators show increased immunosuppressive infiltration at recurrence, while Clearers show decreased infiltration (p < 0.0001 by Kruskal-Wallis).

**Figure 4:**
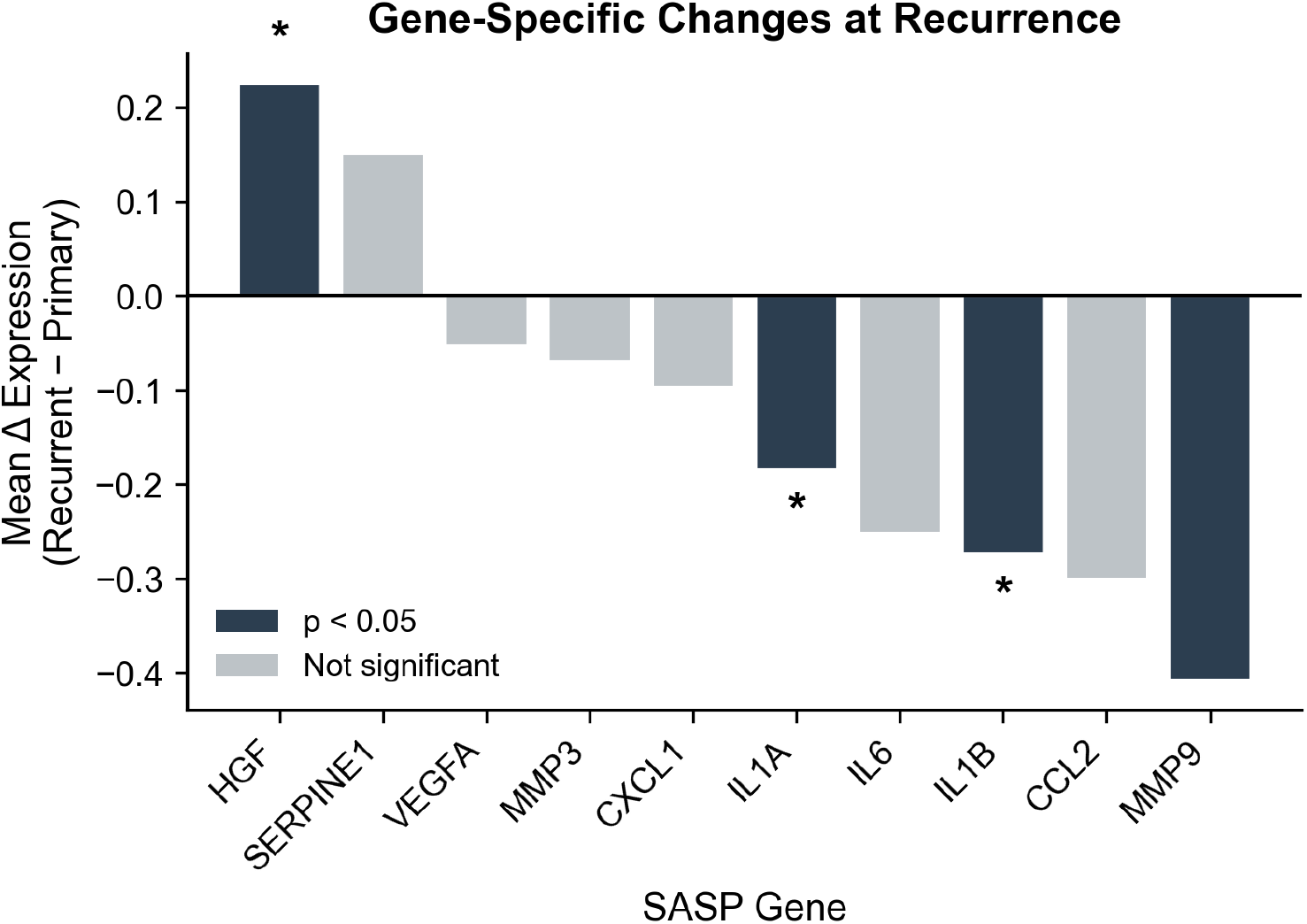
Gene-Specific Changes at Recurrence. (A)Waterfall plot showing mean change in expression for each SASP gene from primary to recurrent tumor. HGF is the only gene that significantly increases (red, p = 0.016). IL1A, IL1B, and MMP9 significantly decrease (blue). Error bars represent standard error. *p < 0.05, **p < 0.01.

These findings are consistent with recent work demonstrating that senescence drives immunotherapy resistance through induction of an immunosuppressive tumor microenvironment^[21]^. Senescent cells evade immune clearance via HLA-E-mediated inhibition of NK and CD8+ T cells^[22]^, and IL-6 secreted as part of the SASP induces CD73 expression on tumor-associated macrophages, creating an adenosine-rich immunosuppressive milieu^[23]^. Recent GBM-specific work has identified a DDX58-STAT1-CSF1 axis in therapy-induced senescent tumor cells that actively recruits tumor-associated macrophages^[24]^.

### Survival Analysis

Baseline SASP score at primary diagnosis was significantly associated with overall survival (Figure S1B). Each 1-unit increase in SASP score was associated with a 65% higher hazard of death (HR = 1.65, 95% CI: 1.34–2.04, p < 0.0001). This effect was driven primarily by Classical subtype tumors (HR = 1.79, p < 0.0001). Trajectory-based survival analysis did not show significant differences between Accumulators, Stable, and Clearers (log-rank p = 0.88; Figure S1A). This is expected given that PMT, rather than survival, is the proximate biological outcome of SASP accumulation, with survival influenced by numerous additional factors including subsequent therapies.

### External Validation in TCGA

The TCGA GBM cohort validated the association between SASP and molecular subtype (Figure 5A). Mesenchymal tumors had significantly higher SASP scores than Classical tumors (Cohen’s d = 1.75, p < 0.0001). While TCGA lacks matched longitudinal data to directly validate ΔSASP, this cross-sectional finding confirms that the SASP-Mesenchymal association identified in GLASS is reproducible in an independent cohort. Effect sizes were consistent across cohorts (Figure 5B), with TCGA showing even larger effects for both the MES vs CL comparison and SASP-M2 correlation.

**Figure 5:**
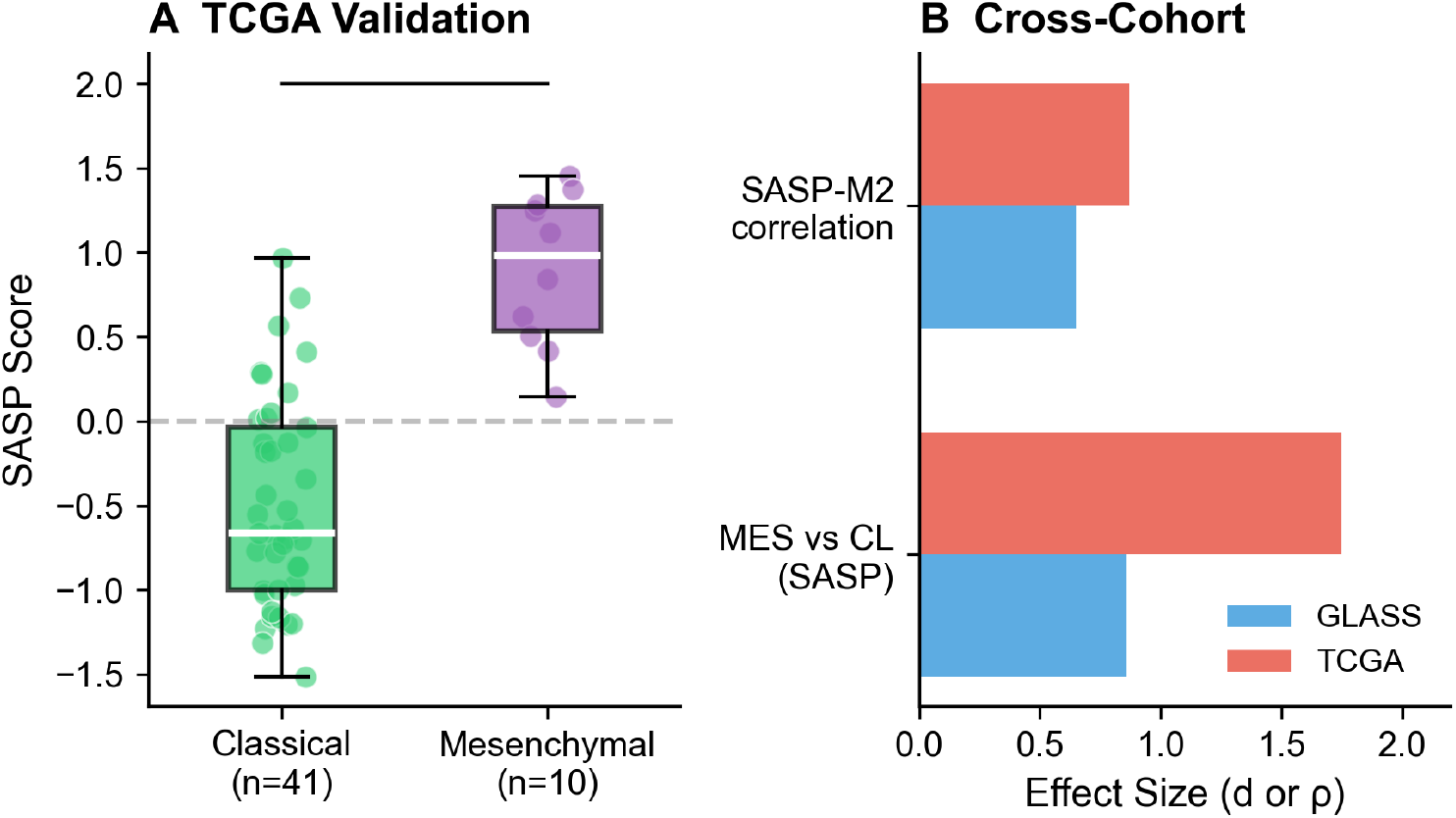
External Validation in TCGA. (A)SASP score by molecular subtype in TCGA GBM cohort (n=52). Mesenchymal tumors have significantly higher SASP than Classical tumors (p < 0.0001, Cohen’s d = 1.75). (B) Cross-cohort comparison of effect sizes. Both the MES vs CL difference and SASP-M2 correlation replicate in TCGA with similar or larger effect sizes.

### External Validation in CGGA

The CGGA cohort (n=225 GBM) provided independent validation of key findings (Figure 6). Predicted Mesenchymal tumors had significantly higher SASP than Classical tumors (Cohen’s d = 1.14, p < 0.0001; Figure 6A), confirming the SASP-subtype association with a large effect size. Baseline SASP predicted worse survival (HR = 1.35, 95% CI: 1.11–1.63, p = 0.003; Figure 6B), and the SASP-M2 macrophage correlation was even stronger than in GLASS (ρ = 0.81, p < 0.0001; Figure 6C). In IDH-wildtype tumors specifically, SASP remained prognostic in both GLASS (HR = 1.32, 95% CI: 1.06–1.66, p = 0.015) and CGGA (HR = 1.35, p = 0.003), confirming the clinical relevance in the most common GBM molecular subtype.

**Figure 6:**
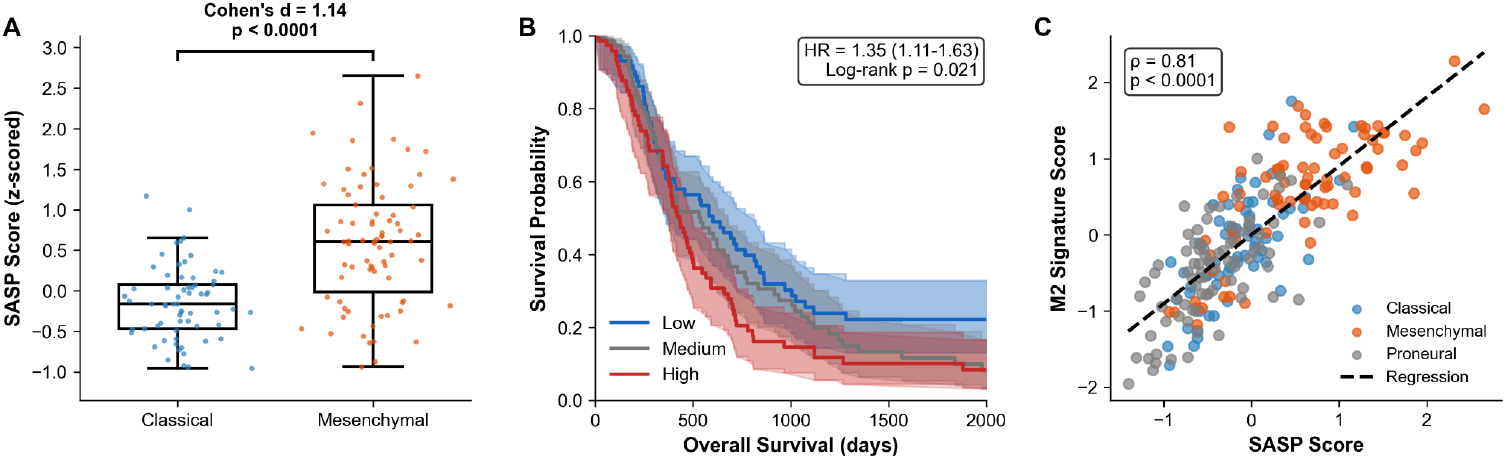
External Validation in CGGA. (A) SASP score by predicted molecular subtype in CGGA GBM cohort (n=225). Mesenchymal tumors have significantly higher SASP than Classical tumors (p < 0.0001, Cohen’s d = 1.14). Subtypes were predicted using Verhaak 2010 classifier signatures. (B) Kaplan-Meier survival curves by SASP tertile. Higher SASP is associated with worse survival (HR = 1.35, 95% CI: 1.11–1.63, log-rank p = 0.021). (C) Correlation between SASP score and M2 macrophage signature. Points colored by predicted subtype. Spearman ρ = 0.81, p < 0.0001.

### Sensitivity and Robustness Analyses

The SASP-survival association was consistent across multiple signature definitions (HR range: 1.32–1.35, all p < 0.01; Supplementary Figure 2C), indicating robustness to gene selection. SASP was distinct from generic inflammation (r = 0.47, below the 0.5 threshold; Supplementary Figure 2A), and SASP retained strong prognostic significance after adjusting for inflammation score in multivariate Cox regression (HR = 1.99, 95% CI: 1.61–2.46, p < 0.0001; Supplementary Figure 2B). The inflammation score itself was not prognostic (HR = 0.93, p = 0.50), confirming that SASP captures senescence-specific biology beyond generic inflammation. ΔSASP was independent of treatment modality (Kruskal-Wallis p = 0.68).

## Discussion

### Summary of Principal Findings

This study provides the first longitudinal analysis of SASP dynamics in matched primary-recurrent glioblastoma pairs. Our key findings are:

1. SASP dynamics are subtype-specific: Mesenchymal tumors show significant SASP clearance (p = 0.0001), while Classical tumors remain stable.
2. ΔSASP strongly predicts PMT: Each unit increase in ΔSASP confers 5-fold higher odds of mesenchymal transition (OR = 5.06).
3. Clearers are protected: Only 2.4% of Clearers undergo PMT, compared to 27.3% of Accumulators.
4. HGF is the exception: While most SASP factors decrease, HGF uniquely increases at recurrence.
5. ΔSASP correlates with immunosuppression: Strong association with M2 macrophage infiltration (ρ = 0.65).

### Biological Interpretation: Why Subtype-Specific Effects?

The striking difference between Mesenchymal (ΔSASP = −0.83) and Classical (ΔSASP = +0.03) tumors may reflect differences in senescence surveillance capacity. Mesenchymal GBMs are characterized by extensive immune infiltration, including high proportions of tumor-associated macrophages^[12,25]^. These immune cells may more effectively clear senescent tumor cells through senescence surveillance mechanisms. Classical tumors, with their lower immune infiltration, may lack this clearance capacity.

When senescent cells do accumulate in Classical tumors, the resulting SASP may drive transition toward a Mesenchymal phenotype—explaining why high ΔSASP predicts PMT. This model suggests that PMT represents, in part, a failure of senescence surveillance.

The mechanistic connection between SASP and PMT is supported by their shared transcriptional regulation. NF-κB serves as the master regulator of both the SASP program and mesenchymal differentiation in GBM^[13,14,26]^. Radiation activates NF-κB via ATM, which then induces both SASP factor secretion and mesenchymal transcription factors including C/EBPβ and STAT3^[27]^. This shared regulatory architecture provides a mechanistic basis for the SASP-PMT correlation we observe.

### The HGF Exception: Stromal vs Tumor Senescence

The unique increase in HGF at recurrence, while other SASP factors decrease, suggests a critical distinction between tumor-intrinsic and stromal senescence. Bulk RNA-seq captures both tumor and microenvironment, and the HGF increase likely reflects secretion by senescent stromal cells, particularly astrocytes, rather than tumor cells. Fletcher-Sananikone et al. demonstrated that irradiated brains upregulate CDKN1A (p21) and SASP factors, with senescent astrocytes secreting HGF that activates MET signaling in glioma cells^[18]^. This finding has been extended by recent work showing that HGF-expressing macrophages activate MET on mesenchymal tumor cells, maintaining the aggressive phenotype^[28]^.

\The therapeutic implication is clear: targeting the HGF/MET axis may be valuable regardless of tumor-intrinsic SASP trajectory. Clinical data support this approach: a phase II trial of onartuzumab (anti-MET) plus bevacizumab demonstrated that patients with HIGH HGF expression had significantly improved progression-free survival (HR 0.46, p = 0.0108)^[29]^, identifying HGF as a predictive biomarker for MET-targeting therapy. Additionally, crizotinib (MET/ALK inhibitor) in combination with temozolomide and radiotherapy has shown acceptable safety in newly diagnosed GBM^[30]^.

### Clinical Implications

#### Patient Stratification for Senolytics

Senolytic therapies including dasatinib plus quercetin (D+Q), navitoclax (ABT-263), and fisetin are entering clinical trials for various indications^[31]^, but patient selection biomarkers are lacking. Our data suggest that ΔSASP trajectory could identify patients most likely to benefit. Accumulators, who fail to clear senescent cells and progress to Mesenchymal phenotype, represent logical candidates for senolytic intervention. Clearers, who demonstrate effective senescence surveillance, may not require such therapy.

Several lines of evidence support the therapeutic potential of senolytics in GBM. Salam et al. demonstrated that ABT-263 improved survival by approximately 30% in mouse GBM models^[8]^. Critically, senescent GBM cells acquire selective BCL-xL dependency, with navitoclax IC50 values significantly lower in senescent versus proliferating cells across multiple GBM cell lines^[32]^. A recent study identified cIAP2 as a novel senolytic target, demonstrating that the SMAC mimetic birinapant prevents recurrent tumor emergence in both patient-derived xenograft and immunocompetent models following radiotherapy^[33]^.

For CNS-directed therapy, drug penetration is a key consideration. Recent clinical data confirm that dasatinib crosses the blood-brain barrier and achieves measurable CSF concentrations in humans^[34]^, supporting D+Q as a viable CNS senolytic strategy. Beltzig et al. systematically screened senolytics against TMZ-induced senescent GBM cells and found that ABT-737, navitoclax, fisetin, and artesunate all showed activity^[35]^.

#### Combination Therapy Rationale

The strong correlation between ΔSASP and immunosuppressive infiltration (ρ = 0.65) suggests that senolytics might synergize with immunotherapy. By eliminating SASP-producing senescent cells, senolytics could reduce M2 macrophage recruitment and create a more favorable immune microenvironment for checkpoint inhibitor efficacy. Recent work has demonstrated that ABT-263 reverses the immunosuppressive myeloid cell profile and restores CD8+ T cell proliferation in tumor-bearing mice^[21]^.

### Limitations

Several limitations should be acknowledged. First, bulk RNA-seq cannot distinguish tumor from stromal senescence, which may explain the divergent behavior of HGF versus other SASP genes. Single-cell RNA-seq would be needed to resolve this question. Second, the GLASS cohort was enriched for Classical subtype (87%), limiting power to detect effects in Proneural tumors. Third, we lack functional validation, we did not treat patient-derived cells with senolytics to confirm that trajectory predicts response. Fourth, trajectory-based survival was not significant (p = 0.88), though this is expected given that PMT is the proximate outcome and survival is influenced by many downstream factors including salvage therapies. Fifth, treatment heterogeneity across GLASS participating institutions may introduce confounding that we cannot fully control, though multivariate adjustment for treatment, age, IDH status, and surgical interval did not substantially alter the ΔSASP–PMT association (adjusted OR = 7.95 vs unadjusted OR = 8.13). Robustness analyses confirmed the finding was stable across model specifications (OR range: 5.57–8.13, all p < 0.01), with no influential outliers identified and cross-validation AUC of 0.82 ± 0.08. Additionally, the SASP-survival association was partially confounded by IDH status in multivariate analysis (adjusted HR = 1.21, p = 0.079 in CGGA), though the effect remained significant when analyzed within IDH-wildtype tumors specifically. MGMT methylation data were missing for 36.8% of patients, though missingness was not associated with ΔSASP or PMT status. The longitudinal ΔSASP findings could not be externally validated due to the lack of matched primary-recurrent pairs in available validation cohorts. Finally, while TCGA and CGGA validated the SASP-subtype association, they cannot directly validate ΔSASP due to their cross-sectional design.

### Future Directions

Key next steps include: (1) single-cell RNA-seq to identify senescent cell populations and their origin (tumor vs stroma); (2) functional validation using patient-derived cells stratified by trajectory and treated with senolytics; (3) prospective validation in clinical trials with senolytic agents; (4) development of liquid biopsy approaches for non-invasive trajectory monitoring; and (5) testing whether trajectory can be therapeutically modified.

## Conclusions

This study reveals that SASP dynamics at glioblastoma recurrence are heterogeneous and subtype-specific rather than uniformly accumulating as previously hypothesized. Mesenchymal tumors demonstrate significant SASP clearance, while Classical tumors show no net change. The critical finding is that patients who fail to clear SASP—the “Accumulators”—undergo mesenchymal transition at dramatically higher rates (27% vs 2%) and develop immunosuppressive microenvironments. ΔSASP trajectory thus defines distinct recurrence phenotypes with direct therapeutic implications. Accumulators represent optimal candidates for senolytic therapy, while Clearers demonstrate effective senescence surveillance that might be therapeutically reinforced. These findings establish a biological rationale for biomarker-guided patient selection in senolytic trials and identify SASP accumulation as a potentially modifiable driver of adverse phenotypic transition in GBM.

## Data Availability

GLASS consortium data are available at synapse.org. TCGA data are available via cBioPortal. Analysis code is available upon request. CGGA data are available at cgga.org.cn Analysis code will be deposited in a public GitHub repository upon publication. In the interim, code is available from the corresponding author upon reasonable request.

## Acknowledgments

We thank the GLASS Consortium and all contributing institutions for making longitudinal glioma data publicly available. GLASS data are available through Synapse (syn17038081) under GLASS data use policies. We acknowledge The Cancer Genome Atlas (TCGA) and the Chinese Glioma Genome Atlas (CGGA) for providing validation cohorts.

## Author Contributions

V.K. and A.T conceived the study, performed all analyses, and wrote the manuscript.

