## Supplemental Figures and Tables for "Subtype-Specific SASP Dynamics Predict Mesenchymal Transition in Recurrent Glioblastoma"

### Supplementary Figures and Tables

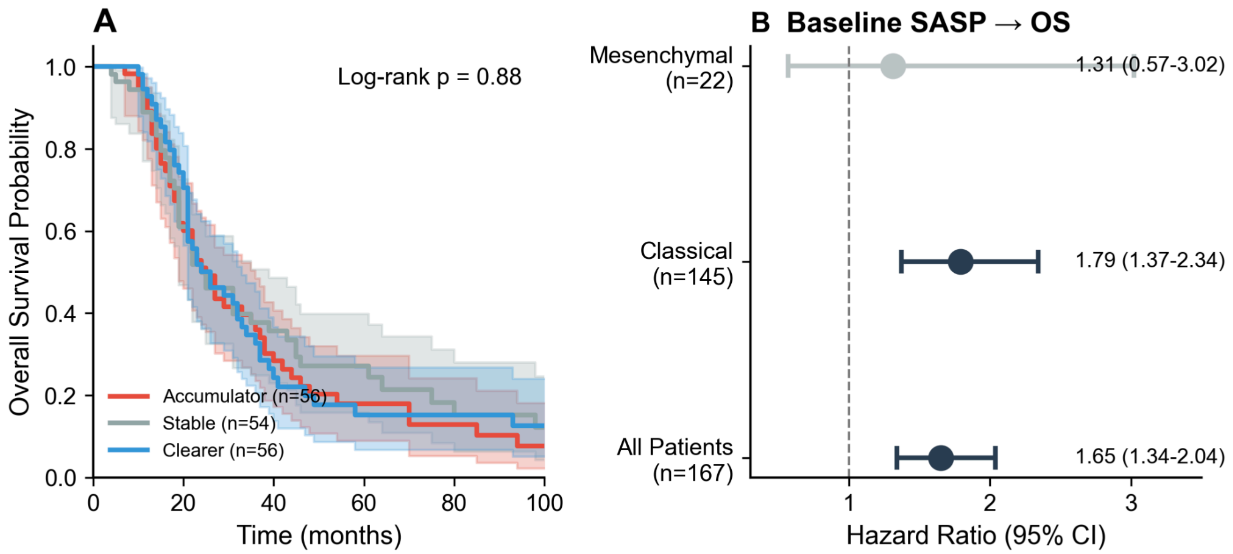


**Supplementary Figure 1: Survival Analysis.**

(A) Kaplan-Meier curves for overall survival stratified by trajectory group. No significant difference between groups (log-rank $p = 0.88$). (B) Forest plot showing hazard ratios for baseline SASP score predicting overall survival. Overall HR = 1.65 (95% CI: 1.34–2.04, $p < 0.0001$). Effect is significant in Classical subtype (HR = 1.79) but not Mesenchymal (HR = 1.31, $p = 0.53$).

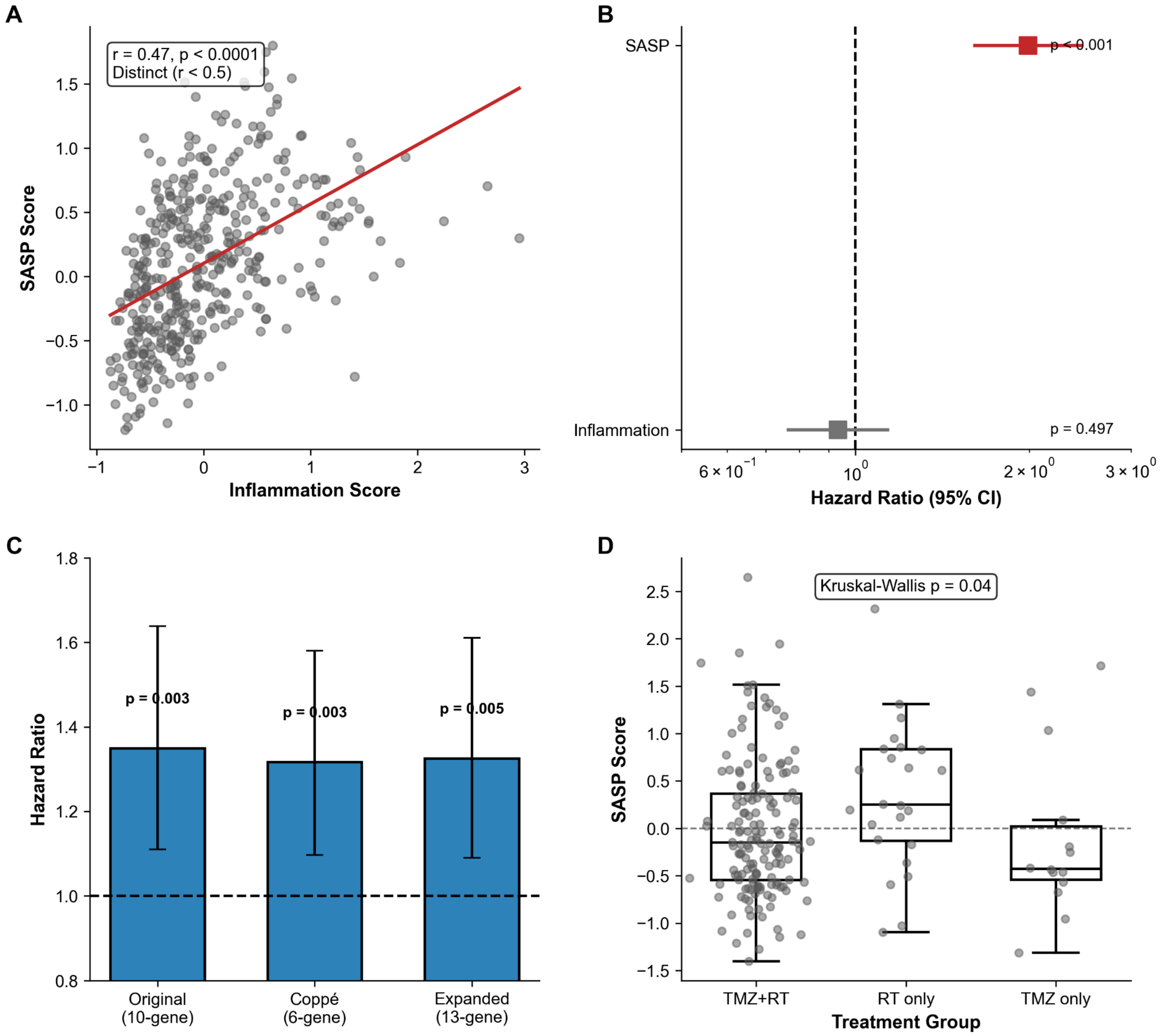


**Supplementary Figure 2: Robustness and Sensitivity Analyses.**
(A) Correlation between SASP score and generic inflammation score (r = 0.47, p < 0.0001). SASP is distinct from inflammation (r < 0.5 threshold). (B) Forest plot of multivariate Cox regression. SASP remains highly significant after adjusting for inflammation (HR = 1.99, p < 0.001), while inflammation score is not prognostic (HR = 0.93, p = 0.50). (C) SASP signature robustness. Hazard ratios are consistent across original 10-gene, Coppé 6-gene, and expanded 13-gene signatures (HR range: 1.32–1.35, all p < 0.01). (D) SASP score by treatment modality (Kruskal-Wallis p = 0.04)

| Gene Symbol | Full Gene Name | SASP Category | Biological Function | Literature Support | Reason for Inclusion/Exclusion |
|---|---|---|---|---|---|

| | | | | | |
|---|---|---|---|---|---|
| IL6 | Interleukin 6 | Pro-inflammatory cytokine | Key pro-inflammatory cytokine that promotes tumor growth, angiogenesis, and immune suppression in GBM | Coppé et al. (2008) *Cell*; Acosta et al. (2013) *Cell* | Included - Canonical SASP factor, highly expressed in senescent cells and GBM microenvironment |
| IL1A | Interleukin 1 alpha | Pro-inflammatory cytokine | Pro-inflammatory cytokine that activates NF-κB signaling and promotes inflammation | Coppé et al. (2008) *Cell*; Acosta et al. (2013) *Cell* | Included - Canonical SASP factor, involved in senescence-associated inflammation |
| IL1B | Interleukin 1 beta | Pro-inflammatory cytokine | Pro-inflammatory cytokine that drives inflammation and immune cell recruitment | Coppé et al. (2008) *Cell*; Acosta et al. (2013) *Cell* | Included - Canonical SASP factor, secreted by senescent cells |
| CCL2 | C-C motif chemokine ligand 2 (MCP-1) | Chemokine | Chemokine that recruits monocytes and macrophages to the tumor microenvironment | Coppé et al. (2008) *Cell*; Krtolica et al. (2001) *PNAS* | Included - Canonical SASP factor, promotes macrophage infiltration in GBM |
| CXCL1 | C-X-C motif chemokine ligand 1 (GRO-α) | Chemokine | Chemokine that recruits neutrophils and promotes angiogenesis | Coppé et al. (2008) *Cell*; Acosta et al. (2013) *Cell* | Included - Canonical SASP factor, involved in immune cell recruitment |
| HGF | Hepatocyte growth factor | Growth factor | Growth factor that promotes cell proliferation, migration, and invasion via MET signaling | Coppé et al. (2008) *Cell*; Birch & Gil (2020) *Nat Rev Cancer* | Included - Canonical SASP factor, promotes GBM invasion and mesenchymal transition |

| | | | | | |
|---|---|---|---|---|---|
| VEGFA | Vascular endothelial growth factor A | Growth factor | Key angiogenic factor that promotes blood vessel formation in tumors | Coppé et al. (2008) *Cell*; Acosta et al. (2013) *Cell* | Included - Canonical SASP factor, critical for GBM angiogenesis |
| MMP3 | Matrix metalloproteinas e 3 | Protease | Extracellular matrix protease that promotes tissue remodeling and invasion | Coppé et al. (2008) *Cell*; Acosta et al. (2013) *Cell* | Included - Canonical SASP factor, promotes ECM degradation and invasion |
| MMP9 | Matrix metalloproteinas e 9 | Protease | Extracellular matrix protease that degrades basement membrane and promotes invasion | Coppé et al. (2008) *Cell*; Acosta et al. (2013) *Cell* | Included - Canonical SASP factor, highly expressed in GBM and promotes invasion |
| SERPINE 1 | Serpin family E member 1 (PAI-1) | Other | Plasminogen activator inhibitor that regulates fibrinolysis and cell migration | Coppé et al. (2008) *Cell*; Kortlever et al. (2006) *PNAS* | Included - Canonical SASP factor, associated with senescence and poor prognosis |
| CXCL8 | Interleukin 8 (IL-8) | Chemokine | Pro-inflammatory chemokine that recruits neutrophils and promotes angiogenesis | Coppé et al. (2008) *Cell*; Acosta et al. (2013) *Cell* | **Excluded** - Canonical SASP factor but not consistently detected in GLASS expression data; 10-gene signature chosen for robustness |

**Supplementary Table 1: SASP Gene Signature Definition and Rationale**

Complete list of genes considered for the SASP signature, including the 10 genes used in the final score calculation and CXCL8 which was excluded due to data unavailability. Genes are categorized by SASP function and supported by canonical senescence literature. All included genes were consistently detected across GLASS, TCGA, and CGGA datasets.

| Characteristic | Accumulator (n=56) | Stable (n=55) | Clearer (n=56) | p-value |
|---|---|---|---|---|
| Age at diagnosis, years* | 51.0 (42.0–64.0) | 49.5 (43.2–60.0) | 51.0 (38.8–56.5) | 0.741 |
| IDH status, % wildtype | 73.6 | 65.2 | 74.4 | 0.558† |
| IDH status, % mutant | 26.4 | 34.8 | 25.6 | |
| Primary subtype, % Classical | 100.0 | 98.0 | 100.0 | 0.362† |
| Treatment, % TMZ | 56.1 | 68.0 | 63.6 | — |
| Treatment, % RT | 52.6 | 54.0 | 54.5 | — |
| Treatment, % both | 35.1 | 36.0 | 38.6 | — |
| Time to recurrence, months* | 11.0 (7.0–24.0) | 11.0 (7.0–18.0) | 12.5 (8.8–22.5) | 0.365 |
| MGMT methylation, % methylated | 36.8 | 32.0 | 38.6 | 0.540† |
| MGMT methylation, % unmethylated | 21.1 | 32.0 | 29.5 | |
| Baseline SASP score* | −0.368 (−0.657–0.095) | 0.054 (−0.606–0.365) | 0.230 (−0.109–0.673) | <0.001 |

**Supplementary Table 2: Baseline Patient and Tumor Characteristics Stratified by ΔSASP Trajectory Group**

Clinical and molecular characteristics of patients classified as Accumulators (n=56), Stable (n=55), or Clearers (n=56) based on tertile cutoffs of ΔSASP. *Continuous variables presented as median (IQR) and compared using Kruskal-Wallis test. †Categorical variables compared using Chi-square test. TMZ, temozolomide; RT, radiotherapy; MGMT, $O^6$-methylguanine-DNA methyltransferase. Groups were well-balanced across baseline clinical characteristics, with the exception of baseline SASP score, which was lowest in Accumulators and highest in Clearers by design due to regression to the mean effects ($p < 0.001$).

**A. Logistic Regression for Proneural-to-Mesenchymal Transition**

| Model | Variable | OR | 95% CI | p-value |
|---|---|---|---|---|
| **Model 1 (Unadjusted)** | | | | |
| | ΔSASP (continuous) | 8.06 | 3.19–20.40 | <0.0001 |
| **Model 2 (Adjusted)** | | | | |

| | | | | |
|---|---|---|---|---|
| | ΔSASP (continuous) | 7.83 | 2.94–20.84 | <0.0001 |
| | Age (continuous, per year) | 1.03 | 0.98–1.08 | 0.246 |
| | IDH status (wildtype vs mutant) | 4.03 | 0.73–22.25 | 0.109 |
| | Time to recurrence (continuous, per month) | 1.00 | 0.96–1.04 | 0.863 |
| | Treatment (both vs other) | 0.81 | 0.25–2.62 | 0.726 |

**B. Cox Proportional Hazards Regression for Overall Survival**

| Model | Variable | HR | 95% CI | p-value |
|---|---|---|---|---|
| **Unadjusted** | | | | |
| | SASP score (continuous) | 1.66 | 1.27–2.16 | 0.0002 |
| **Adjusted** | | | | |
| | SASP score (continuous) | 1.19 | 0.85–1.65 | 0.310 |
| | Age (per year) | 1.03 | 1.01–1.05 | 0.005 |
| | IDH status (wildtype vs mutant) | 2.85 | 1.88–4.31 | <0.0001 |
| | Primary subtype (Mesenchymal vs Classical) | 1.45 | 0.88–2.39 | 0.142 |

**Supplementary Table 3: Multivariate Regression Models for Proneural-to-Mesenchymal Transition and Overall Survival**

**(A)** Logistic regression models predicting proneural-to-mesenchymal transition (PMT) among patients with non-Mesenchymal primary tumors (n=145). Model 1 shows unadjusted association between ΔSASP and PMT. Model 2 adjusts for age, IDH status, time to recurrence, and treatment modality. ΔSASP remained a strong independent predictor of PMT after adjustment (adjusted OR = 7.83, $p < 0.0001$). **(B)** Cox proportional hazards models for overall survival in the full cohort (n=167). Baseline SASP score at primary diagnosis was associated with worse survival in unadjusted analysis (HR = 1.66, $p = 0.0002$), but this effect was attenuated and no longer significant after adjusting for age, IDH status, and primary subtype (adjusted HR = 1.19, $p = 0.310$), suggesting the prognostic effect of SASP is mediated largely through established molecular features. OR, odds ratio; HR, hazard ratio; CI, confidence interval; TMZ, temozolomide; RT, radiotherapy.
